# Mechanically Sheared Axially Swept Light-Sheet Microscopy

**DOI:** 10.1101/2024.04.10.588892

**Authors:** Jinlong Lin, Dushyant Mehra, Zach Marin, Xiaoding Wang, Hazel M. Borges, Qionghua Shen, Seweryn Gałecki, John Haug, Kevin M. Dean

## Abstract

We present a mechanically sheared image acquisition format for upright and open-top light-sheet microscopes that automatically places data in its proper spatial context. This approach, which reduces computational post-processing and eliminates unnecessary interpolation or duplication of the data, is demonstrated on an upright variant of Axially Swept Light-Sheet Microscopy (ASLM) that achieves a field of view, measuring 774 x 435 microns, that is 3.2-fold larger than previous models and a raw and isotropic resolution of ∼420 nm. Combined, we demonstrate the power of this approach by imaging sub-diffraction beads, cleared biological tissues, and expanded specimens.

## 1. Introduction

The study of biological processes within intact tissues has gained paramount importance in modern biology and pathology. Advances in sample preparation methods, such as tissue clearing and expansion microscopy [1, 2], alongside improvements in optical imaging systems, now make it feasible to investigate sub-cellular biological processes in their native tissue environments [3, 4]. This approach is especially valuable. For example, in pathology, volumetric data can reveal critical insights into rare events, like the identification of isolated metastatic breast cancer cells in lymph nodes [5]. Likewise, in cancer biology, the patterns of cancer dissemination and the complexities of the tumor microenvironment are most evident when observed in its three-dimensional entirety, which offers a more comprehensive view of cellular heterogeneity, immune infiltration, and architectural alterations in adjacent tissues [6].

Light-sheet fluorescence microscopy (LSFM) has emerged as a powerful tool for volumetric imaging, offering fast image acquisition speeds and minimal photobleaching. In LSFM, a 3D volume is acquired by illuminating the specimen from the side, and serially imaging adjacent 2D sections within the specimen with a scientific camera. In classical LSFM geometries, one synchronously sweeps the illumination beam and the detection objective, or the specimen, along the optical detection axis (Figure 1a). Alternatively, as is common in lattice light-sheet microscopy [7] open-top [8] LSFMs, the specimen is scanned obliquely relative to the illumination and detection axes (e.g., in the S direction, Figure 1B). In this geometry, the thickness of the specimen is limited by the mechanical working distance of the illumination and detection objectives (dashed lines, Figure 1B). For objectives with sufficiently large working distances, both the scan direction and the direction orthogonal to it (e.g., X) are limited only by the travel range of the stages employed. As such, oblique sample scanning presents a significant advantage by enabling practically unlimited imaging in two dimensions [8-11].

**Fig 1.**
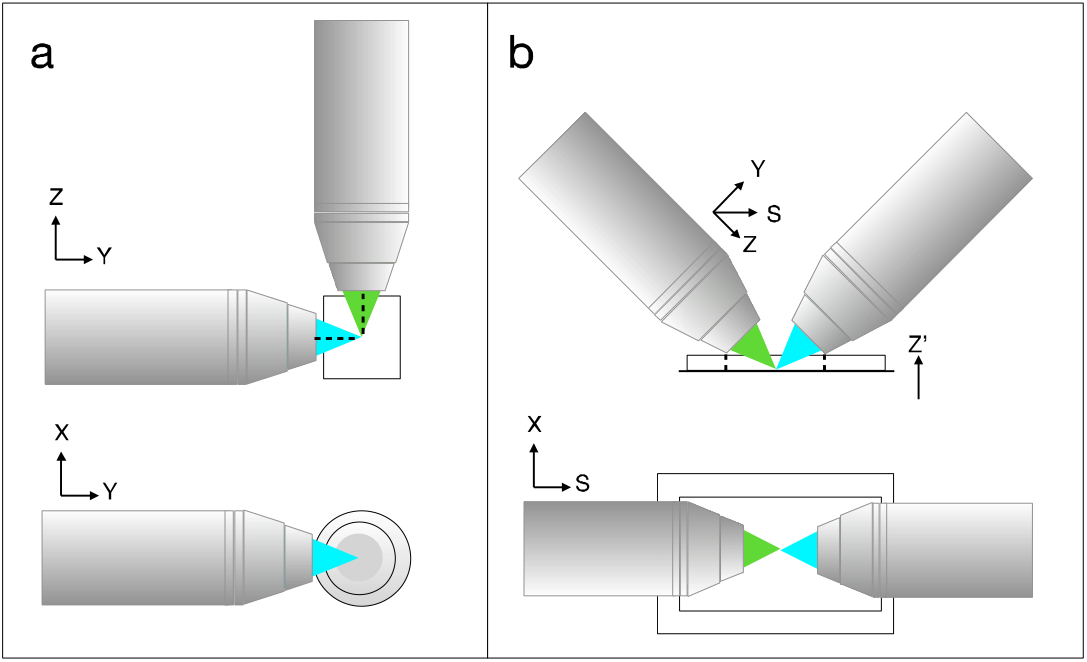
Optical and mechanical constraints on specimen size. (A) When the sample, or laser, is scanned along the detection axis (e.g., Z) to acquire a volume, the optical working distance of the illumination and detection objectives (shown as dashed lines) limit the maximum imaging volume. The orthogonal dimension (e.g., X), is effectively unlimited. (B) When the sample is scanned obliquely (e.g., S) relative to the illumination and detection objectives, the thickness of the specimen is limited by the mechanical working distance of the illumination and detection objectives (shown as dashed lines) in the Z’ dimension. However, both S and the dimension orthogonal to S (e.g., X), are both effectively unlimited in such a geometry, enabling interrogation of thin specimens (e.g., ∼2 mm) with very large lateral extends (e.g., >75 mm).

While oblique scanning offers benefits for samples that have large lateral extents—often seen in tissue-derived specimens—the data acquired requires computational shearing. This process not only leads to data duplication but also introduces empty space into the dataset. Consequently, sheared datasets become significantly larger than the original, raw data. And even with performant CPU and GPU-based software [12], computational shearing of data introduces processing delays and implementation hurdles. To address this challenge, we introduce a multi-axis, or mechanically sheared, image acquisition scheme that eliminates the need for computationally intensive post-processing of the data. This technique is inspired by a recent advance in light-sheet microscopy whereby a high-speed mirror galvanometer within the detection path of an LSFM optically sheared the data in real-time by scanning an otherwise stationary image across the camera [13]. While effective for imaging modalities that produce high-aspect-ratio images, such as those in oblique plane microscopes (OPM) [14], it is less applicable to ASLM, where the entire camera sensor is utilized. Instead, our method mechanically shears the data by simultaneously scanning the sample along both the S and Z’ axes, placing the data correctly in its spatial context from the outset.

## 2. Materials and Methods

### 2.1 Tissue Procurement and Preparation

Animal care was conducted in strict compliance with Institutional Animal Care and Use Committee (IACUC) approved protocols at the University of Texas Southwestern Medical Center. For human specimens, no direct interaction or intervention with human subjects was made for biospecimen collection, and all human tissues were provided and deidentified by someone uninvolved in the study. Detailed methods describing the labeling and clearing of tissues in the supplemental document.

### 2.2 Imaging System and Control

The ASLM presented here is constructed in a dual-inverted selective-plane illumination microscopy-like configuration on a two-tiered vibration isolation system (see Figure S1 for a detailed optical layout) [10, 15]. The bottom tier consists of a 36” x 72” x 18” optical table (Performance Series, TMC) with tuned vibration isolators (UltraDamp Series, TMC) that provide greater dampening at low frequencies. The second tier is assembled on top of the optical table and includes 14-inch vibration-isolating posts (DP14A, Thorlabs) that support a damped 24” x 48” x 4.3” optical breadboard (PG-24-4-ML, Newport). This upper tier serves as the platform for all illumination and detection optics. The specimen stage (FTP-2000, ASI), which is used for sample positioning in Z’, X, and S, is directly mounted on the larger, bottom tier. A detailed summary of the microscope’s construction, and a schematic, is provided in the supplemental document. Our microscope uses navigate (https://github.com/TheDeanLab/navigate) to perform all imaging-related tasks [16]. Stage and filter-wheel operation is performed via serial communication with a Tiger Controller (TG8-BASIC, ASI) equipped with TGCOM, TGFW, and 2x TGDCM2 control cards. Analog and digital tasks are performed with a data acquisition chassis (PXIe-1073, NI) equipped with multifunction input/output and analog output cards (PXIe-6259 and PXI-6733, NI). The acquisition computer (ProEdge SX6800, Colfax International) runs on Microsoft Windows 10 Pro and is powered by an Intel Xeon Silver 4215R CPU @ 3.20 GHz with 96GB of RAM.

### 2.3 Optical Simulations

Optical simulations were performed with Zemax OpticStudio, using manufacturer-provided files or lens specifications. Components were accurately positioned within the virtual model and evaluated with Zemax’s sequential mode.

## 3. Results

### 3.1 Optical Concept

Figure 2 illustrates the fundamental concept of image shearing; when the laser or sample scan axis is not coincident with the optical axes (e.g., as is the case in OPM and diSPIM-like systems), computational shearing of the data is necessary to place it in its proper spatial context (Figure 2a). Shearing of the data is performed in the Fourier domain or with an affine transform, both of which are computationally expensive and laterally shift the data (e.g., in the y-direction) by a factor that depends upon the z-position within the image stack and the angle (α) between the optical and mechanical axes. By laterally shifting the data in a depth-dependent fashion, empty space is introduced into the image canvas, and the overall image size increases.

**Fig 2.**
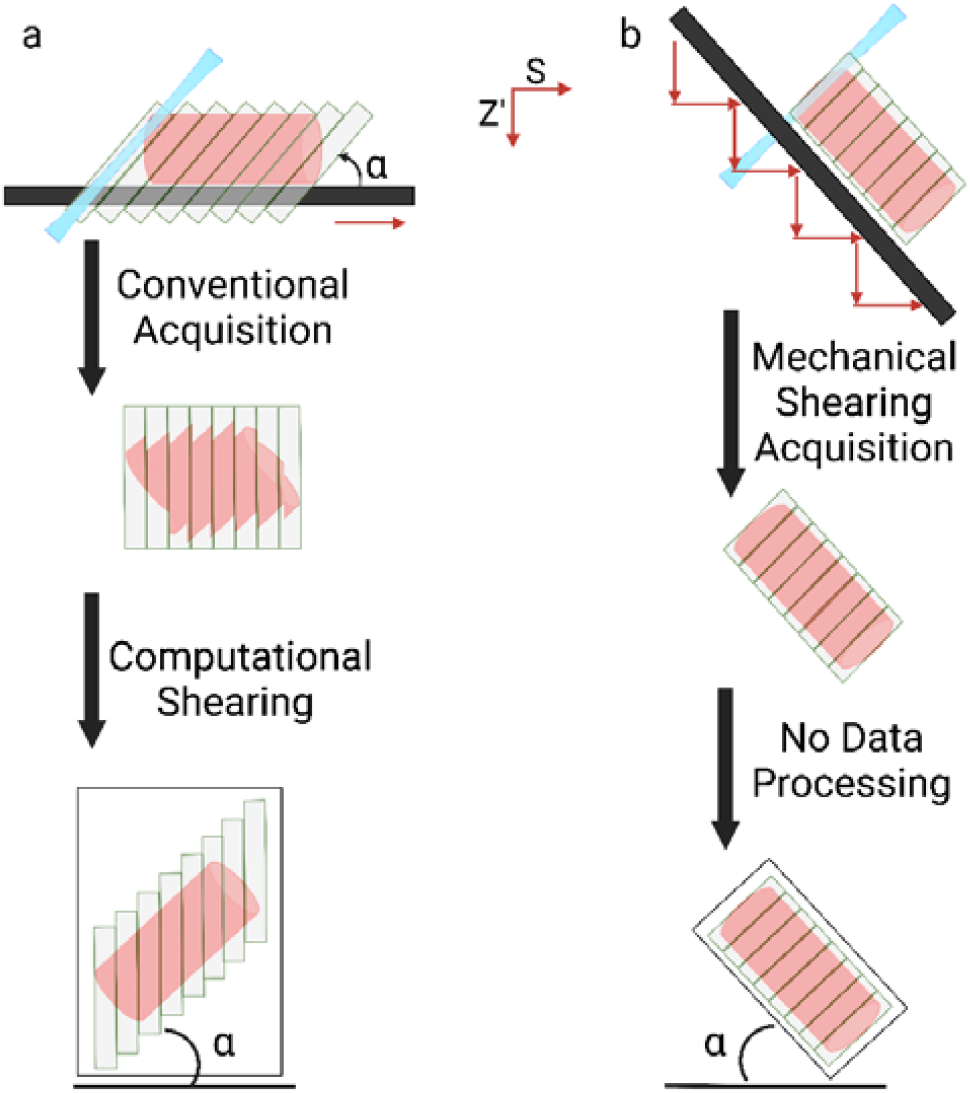
Optical and mechanical shearing of imaging data. (a) In a conventional oblique scan format, the sample is scanned in the S-direction and images are acquired at each adjacent plane in a staggered format (top) but saved in a continuous format (middle). Thus, data must be computationally sheared (bottom) to place it back into its proper spatial context, which introduces empty space above and below the shear axis (see black outline around sheared image). (b) In a mechanically sheared oblique scan format, the sample is simultaneously scanned in S and Z’, thereby placing it in its proper spatial context from the onset. This approach removes excess empty space in the image which significantly reduces memory storage.

Our approach, which we term mechanical shearing, simplifies the imaging workflow by integrating the correction process directly into the data collection stage (Figure 2b). In mechanical shearing, the specimen is simultaneously scanned both vertically and laterally, ensuring that each slice is acquired in its proper orientation from the outset. This method maintains the benefits associated with oblique scanning (e.g., interrogation of thin specimens with large widths and lengths), while avoiding interpolation and thus providing superior resolution. As a result, the imaging process becomes more efficient, reducing both the time and computational resources required to achieve accurately aligned volumetric datasets.

### 3.2 Optical Characterization

To demonstrate the advantages of mechanical shearing for tissue imaging, we developed a microscope in an upright orientation that simultaneously provides a large field of view and high optical resolution. LSFMs face a trade-off between axial resolution and field of view, a limitation observed in LSFMs that adopt both Gaussian and Bessel-Gauss illumination schemes (e.g., lattice light-sheet microscopy[17]). Two notable exceptions to this limitation include dual-view selective plane illumination microscopy (e.g., diSPIM), and Axially Swept Light-Sheet Microscopy (ASLM). In the former, the sample is imaged from orthogonal perspectives, and the data is registered and fused via an iterative deconvolution scheme [15]. In contrast, for ASLM, a diffraction-limited beam is axially scanned synchronously with a camera’s rolling shutter, enabling high-resolution imaging over a large field of view [18]. Data generated has an isotropic resolution and can be viewed in its raw format from any spatial dimension. Thus, we sought to combine the strengths of ASLM and mechanical shearing, making it possible to achieve isotropic imaging in large tissue contexts in an upright microscope geometry without necessitating data manipulation.

To experimentally verify the performance of mechanically sheared data acquired with our upright ASLM, we evaluated the point spread function (PSF) using 200 nm beads embedded in 1% agarose, as depicted in Figure 3a–e. Figure 3a illustrates beads covering the entire camera chip, with zoom-in sections highlighted in Figure 3b. An axial view of the beads is presented in Figure 3c, with three sub-regions displayed in Figure 3d. Figure 3e reveals the PSF of a single 200 nm bead in all three dimensions. To evaluate the spatial uniformity of the resolution, bead images spanning the entire camera chip (774 μm width x 435 μm height) were evenly divided into nine sections. Computer vision routines were utilized to analyze the lateral (XY) Full-Width Half-Maximum (FWHM) of beads within each section (see supplemental document for details). The mean resolution in each section is displayed as a heatmap (Figure 3g), and a slight decrease is observed in the lateral resolution at the image edges compared to the center. To quantitatively assess the isotropy of resolution, data from 200 nm beads spanning the entire camera chip were localized and subjected to a 3D Gaussian fit. The resulting resolution values for all beads in each dimension are plotted in a histogram (Figure 3f) and fit as a mixture of three Gaussian populations. We interpret the Gaussian population with the smallest FWHM as representing single, isolated beads, while Gaussian populations with larger FWHMs are indicative of clusters comprising two or more beads. For each dimension, the largest component of the mixture model was the lower resolution feature, with means of 460, 460, and 483 nm, in X (n=713), Y (n=713), and Z (n=713), respectively. These resolutions are consistent with previously published variants of ASLM, despite a ∼3-fold larger field of view and the mechanically sheared image acquisition format.

**Fig 3.**
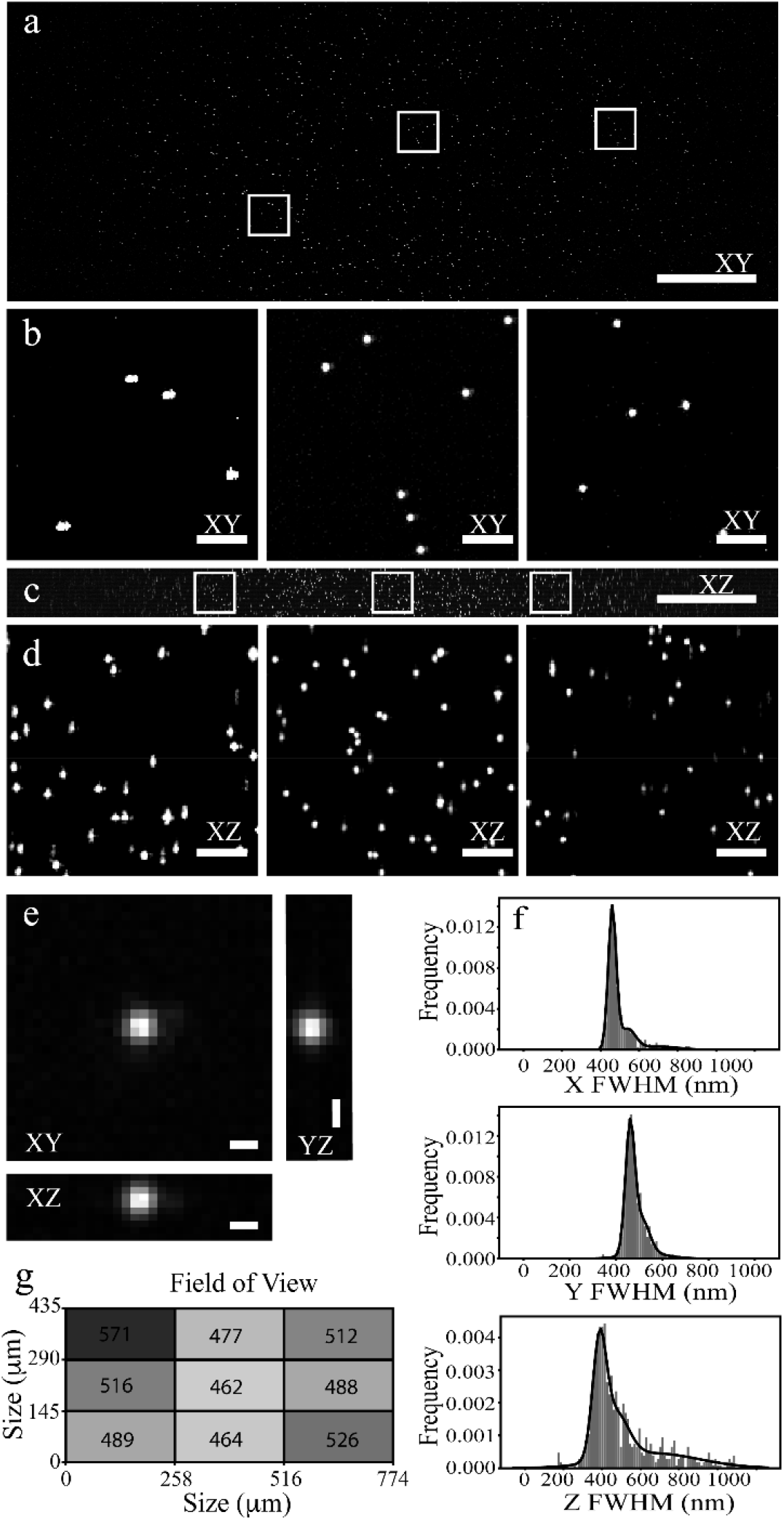
Analysis of 200 nm beads. (a) Displays the XY maximum intensity projection of 200 nm beads in agarose spanning a 20 μm range in the Z dimension. (b) Shows zoomed-in regions of the image from panel a, arranged from left to right. (c) Depicts the XZ maximum intensity projection of 200 nm beads in agarose across a 20 μm range in the Z dimension. (d) Exhibits zoomed-in regions of the image from panel c. (e) Illustrates the maximum intensity projection of a single 200 nm bead in the XY, XZ, and YZ dimensions. (f) Presents histograms of the full-width half maximum (FWHM) of 200 nm beads in the X, Y, and Z dimensions. The number of analyzed beads is 713. (g) Shows a heatmap of the lateral full-width half maximum (FWHM) of 200 nm beads across a 435 μm x 774 μm camera chip. Scale bars: a, c = 100 μm; b, d = 10 μm; e = 1 μm.

#### 3.3 Quantitative Comparison of Computationally and Mechanically Sheared Data

Next, we assessed whether computational shearing, which involves interpolation, exerts a discernible effect on the spatial resolution of a microscope. To evaluate this, 200nm beads were prepared in agarose, and imaged under oblique and mechanically sheared formats. All other imaging variables, including exposure time, z-step size, lateral pixel size, remote focusing amplitude and offset, etc., remained unchanged. Data acquired in the classical oblique scanning format were computationally sheared. Beads from both datasets were evaluated identically using a 3D Gaussian fit, and the FWHMs displayed as a violin plot (Figure 4). Interestingly, the computationally sheared dataset exhibited a slight but statistically significant improvement in lateral resolution (e.g., in X and Y), likely arising from subtle differences in optical alignment. However, a much larger and statistically significant reduction in resolution was observed in the axial dimension for the computationally sheared data. These findings highlight the critical role of interpolation, which effectively functions as a low-pass filter in frequency space, in influencing image resolution.

**Fig 4.**
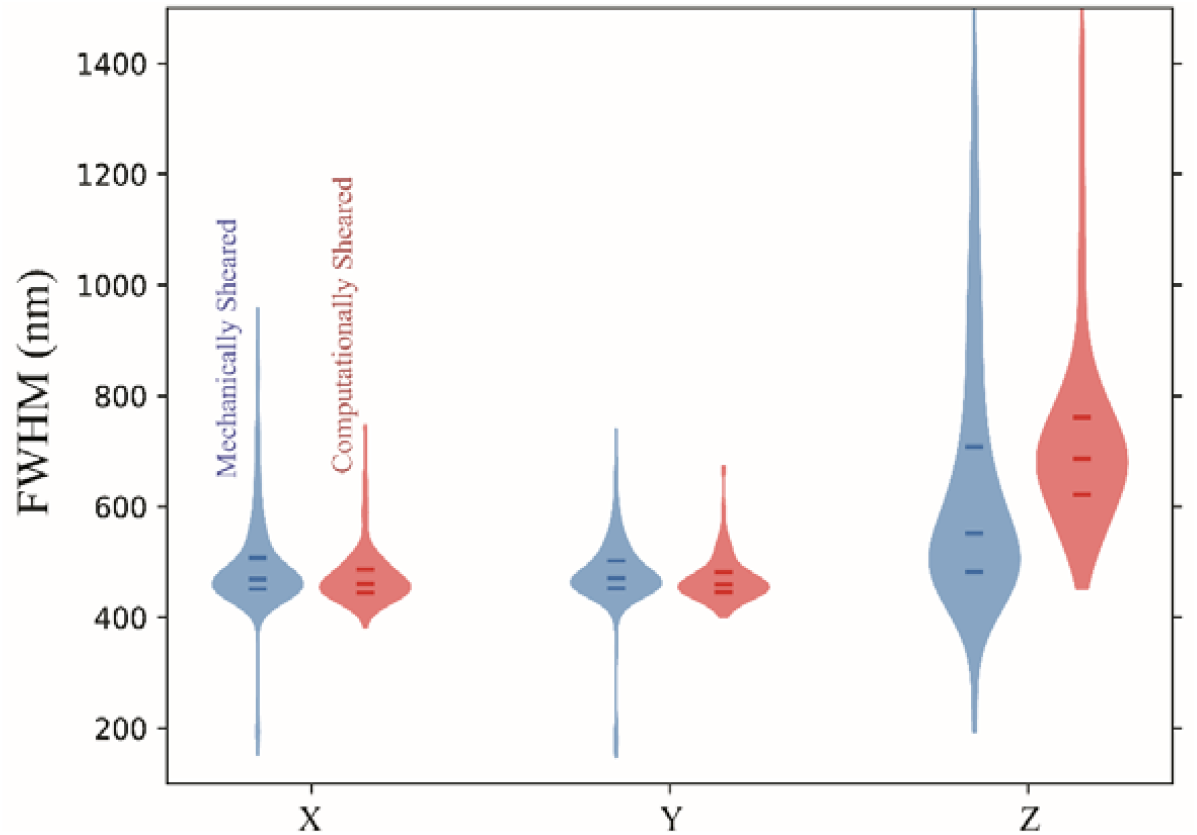
Analysis of 200 nm beads for mechanically and computationally sheared data sets. The mean resolution for mechanically sheared data was 491 nm (X), 477 nm (Y), and 632 nm (Z), whereas for computationally sheared data, it was 473 nm (X), 468 nm (Y), and 723 nm (Z), respectively. Statistical significance was evaluated using both a Welch’s t-test, which assumes that the underlying populations are normally distributed with unequal variances, and a Mann-Whitney U test, which makes no assumptions about the underlying population statistics. All statistical tests were performed with the SciPy toolkit [19].

### 3.4 Overhead Associated with Computational Shearing

Computational shearing of large datasets is associated with significant numerical overhead. And importantly, the sheared data is larger than the input data, which creates additional storage challenges. To evaluate the computational benefits of mechanical versus computational shearing, we conducted benchmarks across a spectrum of CPU and GPU-accelerated computational shearing packages [12, 14, 20]. The outcomes of these benchmarks, detailed in Table 1, include the duration required to shear the data and the dimensions of the sheared output. As anticipated, our analysis revealed that CPU-based methods are only constrained by the available system RAM. Despite this limitation, these methods exhibited considerable processing times (e.g., >20s), even when utilizing advanced file format back-ends designed for distributed, chunked data processing [20]. On the other hand, GPU-accelerated approaches demonstrated a marked improvement in processing speed. However, these methods face limitations due to the GPU’s RAM capacity, typically capped at 24 or 32 GB. This restriction rendered GPU-based methods incapable of processing larger image stacks (e.g., those with greater than 2500 slices), despite their evident acceleration capabilities.

**Table 1.** Comparison of computational shearing software packages.

| FOV (pixels) | Size (GB) |  |  |  | Processing Time (s) |  |  |
| --- | --- | --- | --- | --- | --- | --- | --- |
|  | Original | LLSM5D | Python | CLIJ | LLSM5D | Python | CLIJ |
| 1024x1024x500 | 1.00 | 1.90 | 1.30 | 1.30 | 7.25±1.73 | 144.60±3.66 | 0.97±0.08 |
| 1024x1024x1024 | 2.00 | 3.00 | 3.50 | 3.50 | 19.70±0.96 | 307.00±8.51 | 1.87±0.07 |
| 1024x1024x1300 | 2.50 | 3.50 | 5.00 | 5.00 | 28.56±0.67 | 572.70±32.76 | 2.51±0.12 |
| 1024x1024x1500 | 2.90 | 3.90 | 6.10 | 6.10 | 31.59±3.27 | 776.00±30.32 | 2.86±0.12 |
| 1024x1024x4295 | 3.30 | 4.30 | 7.50 | 7.50 | 36.69±0.53 | 836.80±98.14 | 3.25±0.48 |
| 1024x1024x2000 | 3.90 | 4.80 | 9.60 | 9.60 | 51.34±0.06 | 1328.50±25.19 | 3.92±0.88 |
| 1024x1024x2500 | 4.90 | 5.80 | 13.80 | N/A | 85.46±4.21 | 1935±97.66 | N/A |
| 1024x1024x3000 | 5.90 | 6.70 | 18.70 | N/A | 124.84±9.99 | 2600.50±226.40 | N/A |
| 1024x1024x3500 | 6.80 | 7.70 | 24.40 | N/A | 164.98±0.46 | 3143.75±355.51 | N/A |
| 1024x1024x4295 | 8.40 | 9.20 | 34.80 | N/A | 188.59±13.68 | 5964.00±645.80 | N/A |

### 3.5 Cleared Tissue Imaging

Next, we sought to demonstrate the advantages of mechanical shearing on cleared tissue specimens. Figure 5 displays a human kidney section that was stained with FLARE [21] and cleared with Benzyl Alcohol Benzyl Benzoate (BABB) [22]. Given our large field of view, an entire human nephron could be captured in a single acquisition (Figure 5a). Additionally, a zoomed-in section (Figure 5b & 5c) reveals detailed features within a glomerulus such as individual erythrocytes with their canonical biconcave disc morphology.

**Fig 5.**
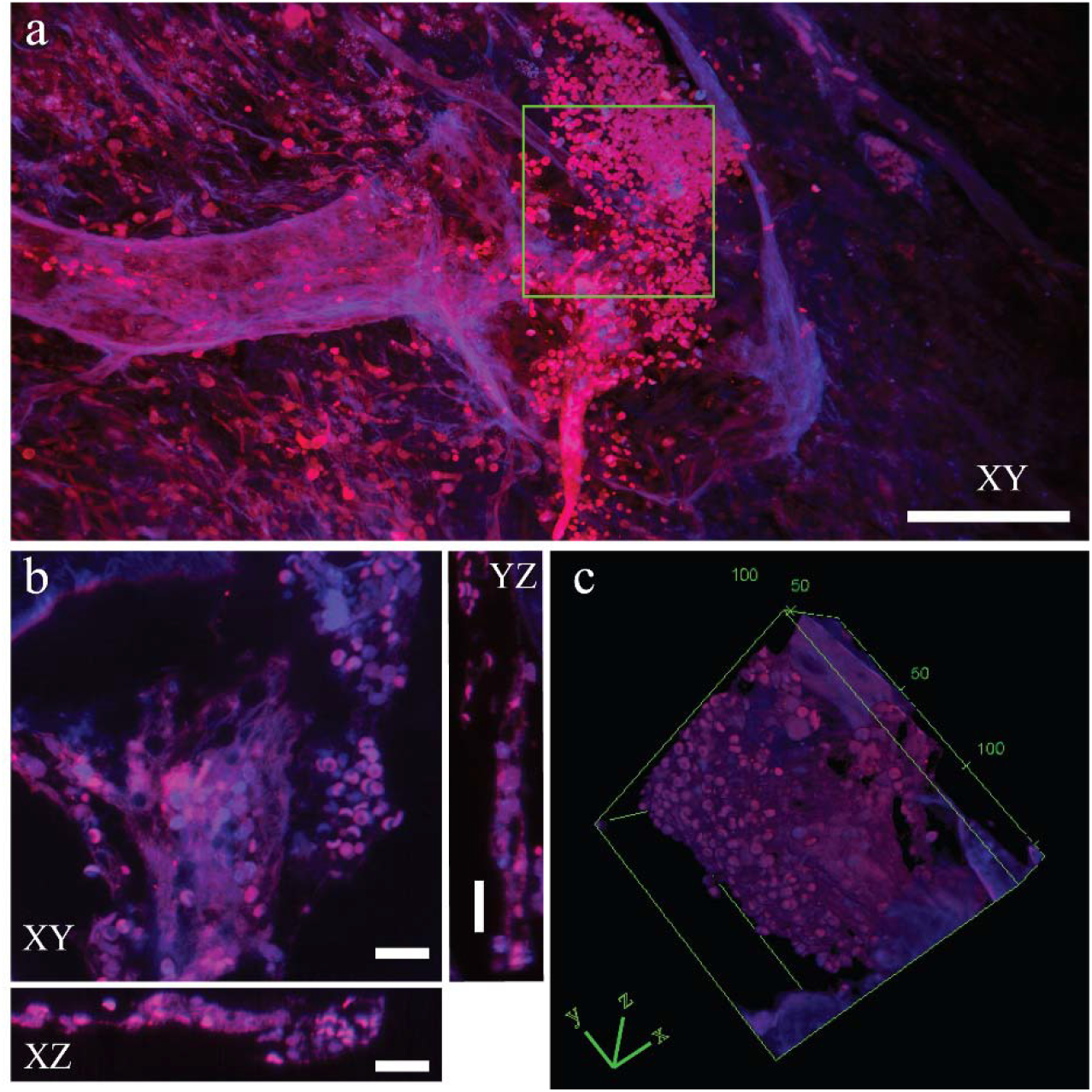
BABB cleared human kidney imaged with mechanical shearing. Specimen was labeled with FLARE [21], and carbohydrates are shown as blue, and proteins are shown as red. (a) Maximum intensity projection of a human nephron. (b) Zoom in a single slice of the region highlighted in image (a). Glomerulus and red blood cells in 3 different dimensions. (c) A volume rendering of glomerulus and red blood cells from (b). Scale bars: a = 100 μm; b, c = 20 μm.

### 3.6 Expanded Tissue Imaging

Expansion microscopy represents a robust approach for imaging biological specimens at sub-diffraction scales. However, unless reinforced with a secondary polymer, expanded tissues are mechanically fragile and thus difficult to image when mounted vertically in a light-sheet microscope. Placing the expanded specimen on a horizontal surface, where it rests under its own weight, avoids the need for secondary embedding of the specimen, simplifying imaging. Moreover, due to the isotropic enlargement of expanded hydrogels, obliquely scanning of the specimen, which provides practically unlimited travel in two of three dimensions, is particularly advantageous. For example, a single coronal mouse brain section, after 4.5-fold expansion, spans 36 mm x 27 mm laterally. To demonstrate the advantages of our mechanically sheared acquisition format for imaging expanded samples, we imaged mouse liver sections. Figure 6(a & b) illustrates the expansion of mouse liver tissue following the protein retention expansion microscopy protocol. In the magnified section depicted in Figure 6(c & d)., the morphology of cancer nuclei is revealed with remarkable detail, including nucleoli. Non-muscle myosin 2A staining, as demonstrated in Figure 6(e & f), highlights clusters of metastatic cells. Figures 6(g, h, and i) showcase mouse liver tissue stained to highlight nuclei and collagen I, a key extracellular matrix component.

**Fig 6.**
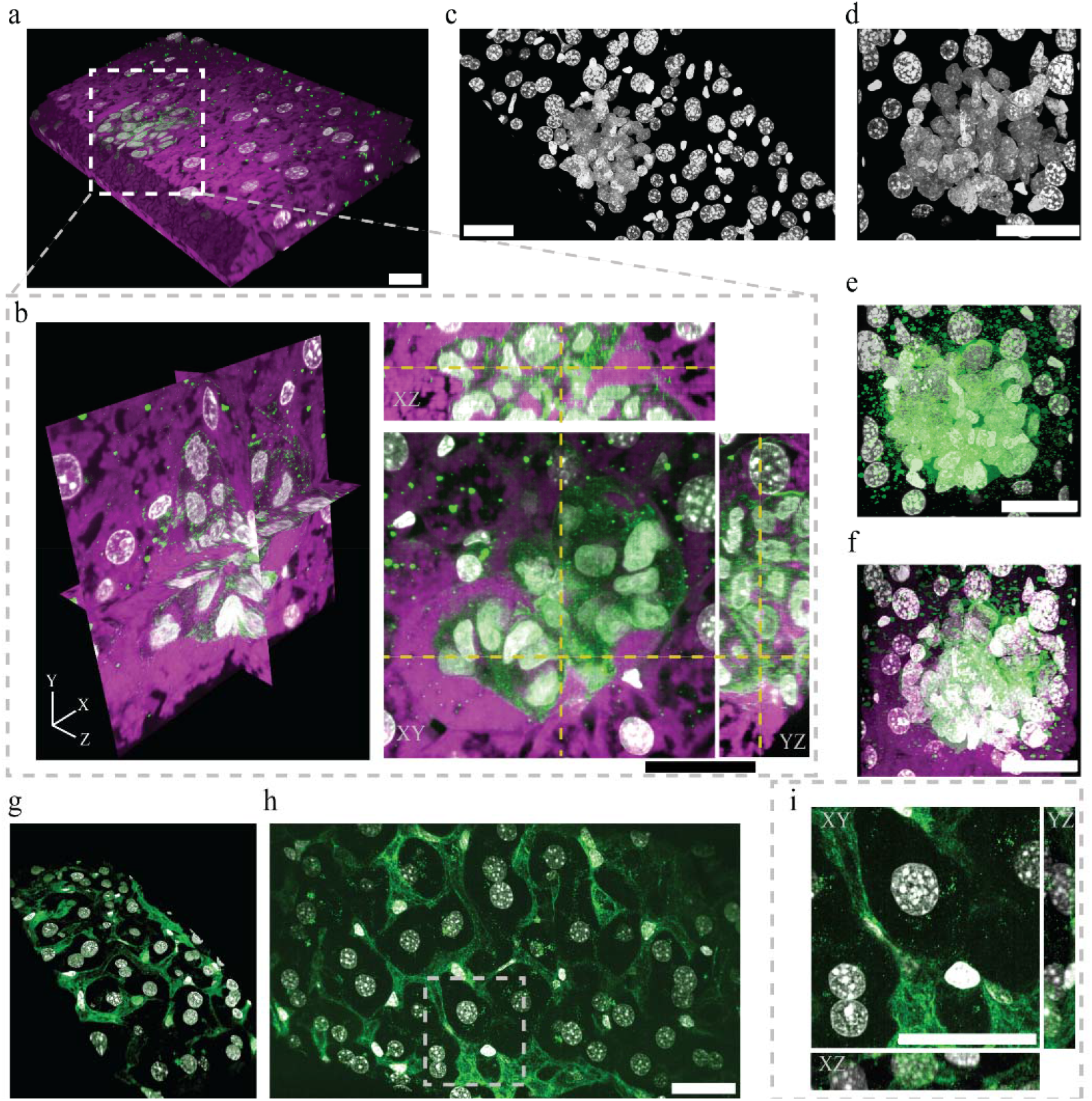
Protein retention expansion microscopy of mouse liver tissues. (a) A three-dimensional volume rendering showcasing mechanically sheared ASLM images of expanded mouse liver tissue exhibiting melanoma micro-metastases. The rendering displays a volume measuring 774.14 x 418.66 x 100 μm. (b) Orthogonal planes from a 287.28 x 261.07 x 100 μm isotropic volume, provide a detailed view of the micro-metastasis. The imaging includes gray for nuclear structures, green for Myosin IIa, and magenta for amines. (c) A 3D projection of the nuclear channel, oriented at a 45-degree angle. (d) A focused view on the melanoma micro-metastasis area of c. (e) Visualization of nuclei with surrounding Myosin IIa signaling in the micro-metastasis region. (f) A merged view incorporating all three channels to illustrate the micro-metastasis area. (g) Volume rendering of M-shearing ALSM images of healthy mouse liver tissue, demonstrating: Gray for nuclear structures and green for Collagen I. (h) A maximum intensity projection offering high-resolution imaging of the sample. (i) Orthogonal planes from the region indicated in h, providing an enhanced view. All images are accompanied by a scale bar measuring 100 μm. The expansion factor was ∼ 4.5 for all samples.

## 4. Discussion

Here, we developed an easy-to-adopt technique termed mechanical shearing that circumvents the need for computationally expensive post-processing of the data. Specifically, the oblique stage geometry permits evaluation of samples with large lateral extents such as clinical specimens and expanded tissues. Once a region of interest is identified, image acquisition proceeds by capturing images after stepping the specimen in both the vertical and lateral dimensions simultaneously. This ensures an accurate spatial representation of the specimen from the outset, thereby avoiding computational shearing, improving resolution, and greatly streamlining the imaging workflow.

To demonstrate the practical benefits of mechanical shearing, particularly for tissue imaging, we constructed an ASLM in an upright, diSPIM-like configuration. To ensure our imaging system’s compatibility with various refractive index solvents and maximize its field of view, we equipped it with high NA multi-immersion objectives and a large format, 12-megapixel CMOS camera. Attempts to maximize the field of view, through integration of a highly corrected 200 mm focal length tube lens resulted in deleterious aberrations at the periphery of the image (data not shown). Thus, guided by Zemax simulations (see Figure S2), we opted for a simple yet effective detection path that included only a 300 mm achromatic doublet. The practical efficacy of our system was validated by measuring the resolution with 200 nm beads embedded in agarose. Importantly, in aqueous solutions, we achieved a spatially uniform and isotropic resolution of ∼460 nm throughout a field of view of 774.14 x 435.46 microns, which is ∼3-fold larger than previous variants. At higher refractive indices, such as BABB, a resolution of ∼330 nm is anticipated.

Owing to the dual-inverted selective-plane illumination microscopy-like geometry, the specimen’s thickness is constrained by the extent to which the objectives’ working distances surpass their physical surfaces, with both objectives at 45 degrees from horizontal and converging on the same focal spot. For NA 0.7 multi-immersion objectives, this amounts to a maximum specimen thickness of 2 mm. However, the other dimensions are limited only by stage travel (here, 120 x 75 mm). While we performed mechanical shearing with stepper motors, an improved configuration would also include a 3D piezo. Such a combination would combine the strengths of large travel range stages with the speed of a piezo and also enable multi-angle projection imaging [13]. Likewise, more complex multi-dimensional stage scanning mechanisms could be used to intelligently adapt the illumination to the contours of the specimen. This adaptability is particularly advantageous for imaging complex biological structures, such as the intricate networks of neuronal tissues or the detailed architecture of vascular systems, where traditional imaging methods may fall short due to their inability to accommodate their unique topographical features. While imaging sensors may be rectangular, the resultant image volumes need not be cuboidal.

By leveraging mechanical shearing in conjunction with a diSPIM-like setup, our approach not only facilitates the exploration of large tissue expanses with enhanced resolution but also significantly reduces the time and computational resources required for post-acquisition processing. Despite the apparent simplicity of multi-dimensional sample scanning (e.g., vector addition), it has surprisingly not been utilized in light-sheet microscopy, as researchers continue to favor more computationally intensive approaches. This methodology eliminates such computational obstacles, thereby reducing the barriers to entry for diverse scientists aiming to understand the molecular origins of tissue function in fields ranging from developmental biology to pathology.

## Acknowledgments

We extend our gratitude to Drs. Reto Fiolka and Gaudenz Danuser for their feedback and insights. Special thanks are due to Dr. Jon Daniels and Steve Saltekoff for their expertise and assistance in optimizing operation of the stages. We also wish to acknowledge Dr. Dana Reed for her continuous support, Dr. Jungsik Noh for his advice on statistical methods, and BioHPC for providing robust computational infrastructure.

## Data availability

Data underlying the results presented in this paper will be made publicly available on Zenodo.

