## Supplemental Document for "Mechanically Sheared Axially Swept Light-Sheet Microscopy"

Mechanically Sheared Axially Swept Light-Sheet Microscopy: supplemental document

1. Materials and Methods

Agarose Beads Sample

A stock solution for the beads was prepared by adding a single drop of 200 nm yellow-green beads (17151-10, Polysciences) to a 50 mL centrifuge tube containing 25 mL of deionized water. The sample was thoroughly mixed with a vortex, and clusters of beads were removed with a 0.22 µm syringe filter (SLGP033RS, Millipore Sigma). To prepare an agarose cube, 50 mg of agarose powder (A9045-25G, Sigma Aldrich) was dissolved into 50 mL of deionized water. The beads were then mixed with the agarose solution in a 2:1 ratio, gently mixed, and placed into a 3D-printed hollow and triangular mold to create a triangular prism-shaped bead specimen. Once the agarose gel had solidified, the bead specimen was placed on a glass slide in such an orientation that the illumination and detection axes were normal to the surface of the triangular prism faces.

Tissue Clearing

A formalin-fixed and paraffin-embedded (FFPE) human kidney (1.2 x 2.3 cm) was heated for 20 minutes at 65°C and then placed in xylene overnight at room temperature to begin deparaffinization. On the next day, the sample was washed in fresh xylene for 3 hours followed by an ethanol gradient (100%, 90%, 80%, 70%) for 3 hours, respectively. After rehydration, the tissue was washed with 1x phosphate-buffered saline (PBS) 3x for 2 hours. To decolorize the tissue, the sample was immersed in 25% QUADROL (122262, Sigma Aldrich) at 37˚C overnight and was refreshed until the supernatant was visibly clear before proceeding with staining. Carbohydrate and amine functional groups were covalently labeled as described in FLARE [1]. After staining, the tissue was dehydrated in a methanol gradient (25%, 50%, 75%, for 1 hour and 100% 2x for 45 minutes), and delipidated 2x for 45 minutes in dichloromethane (DCM) (270997, Sigma Aldrich), ensuring tissues sink before clearing. The final clearing was achieved with repeated fresh Benzyl Alcohol (108006, Sigma Aldrich) and Benzyl Benzoate (BABB, 1:2), (105860010, Thermo Scientific) incubations. After overnight BABB incubation, the sample was ready for high-resolution imaging, having attained optimal transparency through this process.

Expansion Microscopy

All samples were labeled with covalently reactive dyes or via indirect immunofluorescence before hydrogel embedding and expansion using previously established protocols [1-3]. Specifically, a non-perfused and fixed mouse liver was embedded in 4% (w/v) agarose prepared with 1x PBS and sectioned with a vibratome (VT 1000 S, Leica) into 100 μm thick slices. Sections were stored in 1x PBS with 0.02% (w/v) sodium azide at 4°C until use. Tissue sections were then permeabilized and blocked for 6 hours by incubation in Blocking Buffer [0.5% NP40, 10% DMSO, 5% Normal Donkey Serum, 0.5% Triton X-100, in PBS] at room temperature, immunostained with primary antibodies (anti-Myosin IIa (Rabbit mAb #49349, Cell Signaling) or anti-collagen I (MA1-26771, Invitrogen), both at a dilution of 1:100, v/v) overnight at room temperature. This was followed by application of secondary antibodies at approximately 10 µg/mL concentration. For amine staining, samples were incubated for 6 hours with 5 μg/mL ATTO 647N N-hydroxysuccinimidyl ester (18373-1MG-F, Millipore Sigma) in MES buffer.

After staining, the samples were anchored with 0.1 mg/ml Acryloyl-X, SE (Invitrogen) in PBS overnight. Subsequently, they were immersed in a monomer solution [comprising 20% (wt/wt) Acrylamide, 10% (wt/v) Sodium Acrylate, 0.05% (wt/wt) Bis-Acrylamide, and 4% (v/v) Paraformaldehyde] at 4°C overnight. Gelation was induced using 0.2% (wt/v) Ammonium persulfate (APS) and 0.67% (v/v) Tetramethylethylenediamine (TEMED) at 37°C for at least 1 hour. Following gel polymerization, enzymatic digestion (for Figures 5a-f) or heat denaturation (for Figure 5g-i) was performed. Enzymatic digestion involved treating the samples with 8 U/mL Proteinase K (P4850, Millipore Sigma) in Digestion Buffer [200 mM SDS, 200 mM NaCl, and 50 mM Tris base (pH 9.3)] at 37°C for 6 hours. Heat denaturation involved heating the hydrogels at 75 and 90°C for 24 hours each, with incubation in 10mL of Denaturing Solution, consisting of 200mM SDS, 200mM NaCl, 50mM Tris-HCl (pH 9.0), 1x PBS, and deionized water. Post-digestion, the samples were washed in 1x PBS for 15 minutes and stained with SYTOX-Green (Fig. 5a-f), or SYTOX-Orange (Fig. 5g-i) nuclear dyes (1:3000, 1:600, respectively), Invitrogen) in PBS for 1-2 hours. Finally, transparent hydrogels underwent immersion in diH_2_O (10-20 mL) with 3 water exchanges every 20 minutes, ensuring complete physical expansion. Expanded samples were gently affixed to a poly-L-lysine-coated coverslip and imaged promptly.

Optical Layout

The microscope's illumination path begins with laser light originating from an Omicron LightHUB Ultra equipped with lasers emitting at 405, 488, 561, and 642 nm (120, 200, 150, and 140 mW in power, respectively). Upon exiting the fiber, the laser light is collimated with an achromatic doublet lens (AC254-100-A-ML, Thorlabs). It then sequentially passes through an adjustable iris (CP20D, Thorlabs) and its linear polarization orientation is adjusted with a half-wave plate (10RP52-1B, Newport) mounted within a rotation stage (CRM1T, Thorlabs). To form the light-sheet, the collimated laser light is focused with a cylindrical lens (LJ1695RM-A, Thorlabs) onto the surface of a resonant galvanometer (CRS4KHz, Novanta), and recollimated with an achromatic doublet (AC254-75-A-ML, Thorlabs). The orientation of the light-sheet is controlled with a rotation mount (CRM1T, Thorlabs), and the numerical aperture of the light-sheet is adjusted with a mechanical slit (VA100CP, Thorlabs) positioned at the back focal plane of the cylindrical lens. Operation of the resonant galvanometer, which is conjugated to the specimen, results in pivoting of the light-sheet and a reduction in shadow artifacts that arise from optical absorption and/or scattering events [4]. The light is then relayed with a 4f telescope consisting of 150 mm (AC254-150-A-ML, Thorlabs) and 100 mm (AC254-100-A-ML, Thorlabs) achromatic doublets, traverses a polarizing beam splitter (CCM1-PBS251, Thorlabs), a rotation stage (CRM1T, Thorlabs) mounted quarter-wave plate (AQWP3, Bolder Vision), and enters the back pupil of an air objective (20x NA 0.7, LWD S Plan Fl, Nikon Instruments). The beam is focused by the objective onto a pneumatically actuated voice coil that was customized to include a low-mass adaptor and a mirror (LFA-2010, Equipment Solutions). The back-reflected light from the mirror is captured by the same objective, transmitted through the quarter-wave plate, deflected by the polarizing beam splitter, and relayed with a 4f telescope consisting of 125 mm (AC254-125-A-ML, Thorlabs) and 75 mm (AC254-75-A-ML, Thorlabs) achromatic doublets, followed with another 4f telescope consisting of 75 mm (AC254-75-A-ML, Thorlabs) and 150 mm (AC254-150-A-ML) achromatic doublets to the back pupil plane of an NA 0.7 multi-immersion objective (54-12-8, ASI), which illuminates the specimen.

Fluorescence emitted by the specimen is captured orthogonally using an identical NA 0.7 multi-immersion objective (54-12-8, ASI). This light is then focused by a 300 mm achromatic doublet lens (AC508-300-A-ML), transmitted through a 32 mm diameter 8-position filter wheel (FW-1000, ASI) containing four single-band bandpass filters: 442/42 nm (FF01-442/42-32, Semrock), 515/30 nm (FF01-515/30-32, Semrock), 595/31 nm (FF01-595/31-32, Semrock), and 670/30 nm (FF01-670/30-32, Semrock), and detected by a high-speed, scientific CMOS camera (ORCA-Lightning, Hamamatsu) with 4608 x 2592 pixels.

Field Of View Maximization

To maximize the field of view of our imaging system, and enable operation in diverse refractive index solvents, we built an ASLM in an upright configuration with NA 0.7 multi-immersion objectives (54-12-8, ASI) and a large format, 11.6-megapixel scientific CMOS camera (Hamamatsu Lightning, 25.344 x 14.256 mm). Given this detection objective, which has a nominal field of view and focal length of 1 and 8.4 mm, respectively, integration of a 300 mm achromatic doublet provides a magnification of ~29X and aligns the objective's FOV with the camera sensor diagonal (29.078 mm). However, we were uncertain if such a simple lens could provide the requisite corrections over such a large field of view. We thus evaluated aberrations arising from the detection objective before, and after integration of 300 mm achromatic doublet, by modeling the optical system in Zemax’s sequential mode (Figure S2). Despite additional wavefront distortion arising from the achromatic doublet, simulations demonstrated that such a simple detection path could provide a high level of achromatic performance (e.g., with a wavefront error less than 0.125) for a field of view as large as ~800 microns in diameter.

Resolution Measurements

Resolution was assessed using Full-Width Half-Maximum (FWHM) measurements. A resolution analysis pipeline was developed using MATLAB. Initially, beads were identified through thresholding, and their centroids were determined using regionprops3. Beads that touched the image edges and those with a coefficient of determination (R^2^) below 0.9 were excluded. The line profiles of the remaining beads along the X, Y, and Z axes were extracted and fitted with Gaussian functions. The resulting sigma values were converted to FWHM measurements. Subsequently, these FWHM values were exported to CSV files for further analysis using Python.

Computational Benchmarking of Shearing Software

All benchmarking was completed using BioHPC, UT Southwestern’s high-performance computing infrastructure. Tests were performed on a Linux node equipped with an NVIDIA V100 GPU with 32 GB GPU RAM, 512 CPU RAM, and an Intel Xeon Gold 6140 2.30 GHZ CPU processor. For CLIJ benchmarking [5], individual files were loaded into Fiji and a macro was used to record the amount of time required to shear the input image stack using the 3D affine transform function with a shear factor that is defined by the acquisition angle (α=45), mechanical step size (200 nm), and pixel size (143 nm). For LLSM5D benchmarking [6], a MATLAB script was written that loads the input file path into the “crop_deskew_rotate” function and records the time it takes to shear the image according to the input step size, acquisition angle, and pixel size. Mean and standard deviation from 10 consecutive timed runs of CLIJ and LLSM5D were reported. The Python shearing code is a custom software written in our lab which shears the individual images in an input stack by shifting the image in Fourier space by using a Fourier transform according to a shear factor that is defined by the input acquisition angle, step size, and pixel size [7]. The time required to shear the image stack was recorded, and the mean and standard deviation from 5 consecutive timed runs were reported.

2. Supplementary Figures

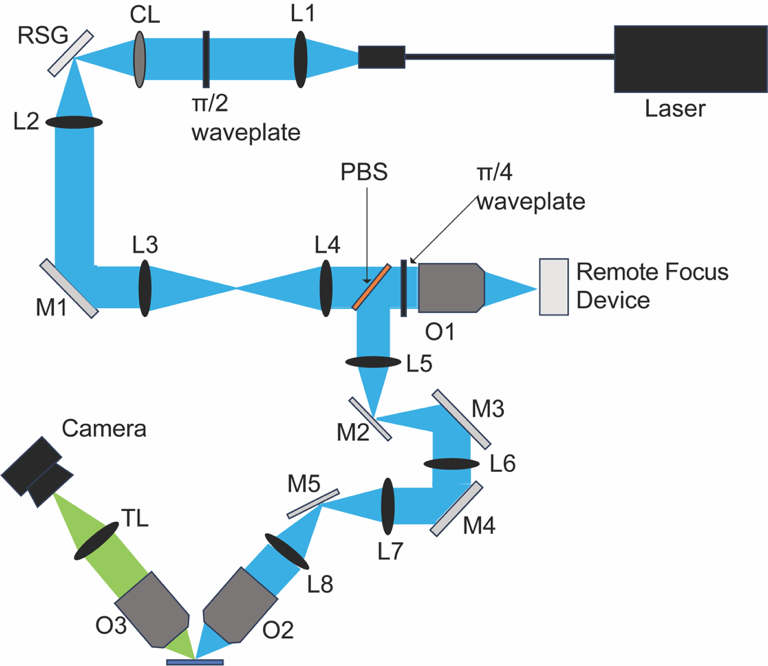

Fig. S1. Optical Layout of Upright ASLM system. L1:L8, achromatic doublet lenses; TL, tube lens; CL, cylindrical lens; RSG, resonant galvo; M1-5, mirrors; π/2 and π/4, achromatic half waveplate, and achromatic quarter waveplate, respectively; PBS, polarized beam splitter; O1, 2, 3, 20X air objective and 2 NA 0.7 multi-immersion objectives, respectively; remote focus device, pneumatically actuated voice coil with mirror; camera, sCMOS Hamamatsu Orca Lightning.

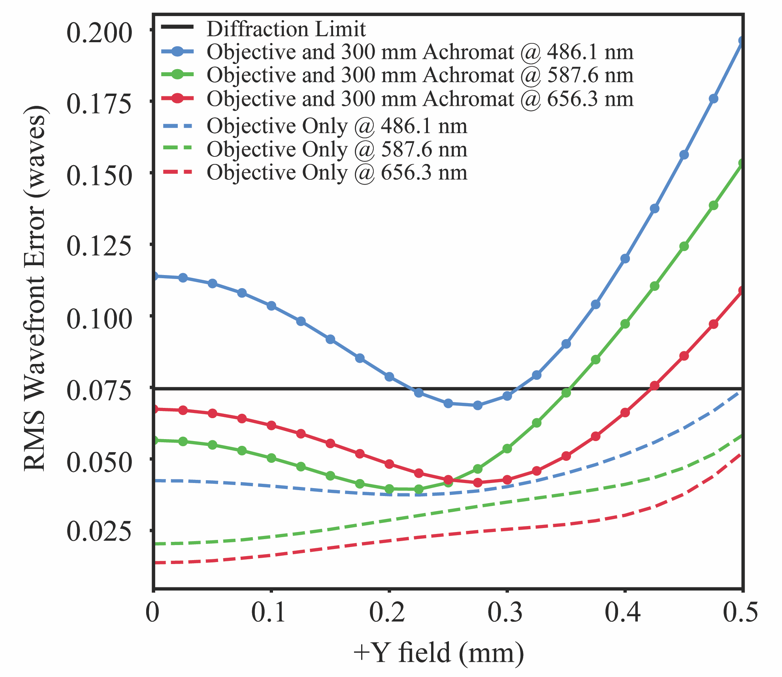

Fig. S2. Optical aberrations for proposed imaging system. Root Mean Square (RMS) wavefront error against the field, with a maximum value of +0.5 mm representing the initial 1 mm FOV of the multi-immersion objective. The dashed lines indicate the wavefront error solely for the objective lens at three different wavelengths, staying below the diffraction limit (represented by the solid black line) across the entire FOV. The solid-colored lines depict the wavefront error post-inclusion of the achromatic doublet. While the error increases across all wavelengths due to the doublet's incorporation, it remains within acceptable limits for our targeted FOV.
